# A streamlined and comprehensive protocol for the generation and multi-omic analysis of human monocyte-derived macrophages

**DOI:** 10.1101/2025.05.30.655544

**Authors:** Olivia G. Palmer, Laurent Perreard, Fred W. Kolling, Patricia A. Pioli, Brittany A. Goods

## Abstract

Macrophages serve both as a first line of defense against invading pathogens and mediate tissue homeostasis. These cells are inherently responsive and heterogeneous and lie on a spectrum of activation states book-ended by M1-like (inflammatory) and M2-like (anti-inflammatory) extremes. The study of human macrophages is necessary to unravel the complex signals and environmental cues that these cells integrate to create their varied phenotypes. *In vitro* protocols to differentiate human monocytes into macrophages use many distinct activation stimuli at variable concentrations and for differing durations of treatment that can impact macrophage fate. These variations can make it challenging to reproduce findings and compare datasets across research environments. Additionally, few protocols to date have performed rigorous characterization with input material from frozen peripheral blood mononuclear cells (PBMCs). This is important since the availability of fresh blood can often be limiting and can lead to a loss of standardized procedures, particularly for cell therapy applications. Here, we have developed a comprehensive protocol to generate human macrophages from monocytes where we rigorously characterize the impact of differentiation conditions and polarization conditions on human macrophages. We provide a detailed protocol for their characterization using several ‘omics readouts, including their cytokine production and transcriptomes. We also perform depolarization experiments to determine durability of macrophage immunophenotype post-removal of polarizing stimuli. Finally, we demonstrate that nuclei can be isolated and profiled by snRNA-seq directly from macrophages in culture, alleviating the need to detach these adherent cells for downstream multi-ome analyses. Taken together, we provide a comprehensive, detailed and streamlined procedure for the differentiation and characterization of human macrophages from monocytes isolated from frozen PBMCs. This is important for enabling the study of macrophages in a more systematic way from biobanked material.

## Introduction

Macrophages are highly plastic immune cells that play critical roles in human health and disease^1,2^. These cells exhibit remarkable functional diversity, contributing to homeostasis, host defense, and tissue repair. Their phenotypic and functional states are shaped by innate and environmental cues, enabling them to adopt a spectrum of activation states that orchestrate nuanced and finely-tuned immune responses and homeostatic functions in tissues^3,4^. Given their essential and pleiotropic roles, understanding the factors that drive macrophage differentiation and activation is crucial for advancing therapeutic strategies targeting inflammatory diseases, infections, healthy aging, and cancer ^5–7^.

Macrophages have been historically classified into broad categories based on their activation state: M0 naïve non-activated macrophages, classically activated M1-like macrophages, which promote inflammatory responses, and alternatively activated M2-like macrophages, which contribute to tissue repair and immune resolution^8–10^. M1-like macrophages, activated by lipopolysaccharide (LPS) and interferon-gamma (IFNγ), typically have elevated expression of surface co-receptors that mediate antigen presentation, such as HLA-DR, CD40, CD80, and CD86, upregulate pro-inflammatory genes including *IDO1, CCR7*, and *GBP1*, and secrete cytokines such as TNF, IL-6, and IL-1β^11–14^. These cells are also characterized by activation of transcription factors like *STAT1*. In contrast, M2-like macrophages, stimulated by IL-4 and/or IL-13, are characterized by high surface markers CD163, CD206 (MRC1), CD209, and the IL-10 receptor, upregulate genes associated with tissue repair and the resolution of inflammation including *ALOX15, F13A1*, and *CCL13*, and secrete anti-inflammatory cytokines such as CCL^17^ and CCL18. Their polarization is driven by transcription factors such as *STAT6*^11,12,15–18^. While this classification provides a useful framework for characterizing cell differences, macrophage activation states exist along a continuum rather than as discrete subtypes, as *in vivo* macrophage activation is derived from local tissue micro-environmental milieux. However, these myriad activation states are bookended by M1-like and M2-like phenotypes^12,16,19^.

The maintenance of macrophage phenotypes *ex vivo* is essential to support cell therapy approaches, mechanistic models with human cells, and for applications in regenerative medicine^20–22^. Despite these important applications, variability in experimental conditions, including cell isolation and culture methods, cytokine concentrations, and donor-to-donor differences, further complicates the ability to define and reproduce macrophage phenotypes *in vitro*^9,10,23,24^. Current methodologies for studying human macrophages often require fresh blood donations, rely on immortalized cell lines, exhibit variability in macrophage yield and activation, and lack standardized protocols, making reproducibility a challenge^25–27^. Finally, few studies have investigated sex differences in *in vitro*-derived macrophages28.

To address these limitations, we optimized an *in vitro* culture system using cryopreserved human monocytes isolated from peripheral blood mononuclear cells (PMBCs) to generate and characterize macrophage activation states under controlled conditions. We evaluated the impact of different cytokine combinations and concentrations on macrophage phenotype using a multi-omic approach. We measured cytokine secretion profiles and performed transcriptomic analysis to uncover the key gene expression signatures that define each activation state. Additionally, we investigated macrophage plasticity by assessing their ability to rapidly depolarize following removal of activation stimuli. Finally, we developed a streamlined workflow to isolate macrophage nuclei for single-nuclei RNA sequencing (snRNA-seq), providing a robust approach for profiling macrophage heterogeneity without the need for harsh dissociation protocols that can rapidly change phenotypes. In the following sections, we present a detailed analysis of macrophage polarization, transcriptional profiles, and functional stability, offering insights into their plasticity, with implications for more standardized *in vitro* modeling for functional studies and cell therapy approaches.

## Results

### Quantifying the impact of various activation stimuli on macrophage phenotype

First, we sought to optimize the differentiation of monocytes into macrophages. Human monocytes were isolated from cryopreserved PBMCs using immunomagnetic isolation of human CD14+ cells and differentiated into macrophages with recombinant macrophage colony stimulating factor (M-CSF). A dose response study was performed with cytokine from different vendors to identify potential differences in baseline activation of these macrophages during differentiation (**Figure S1A**). We observed that that there was very little impact of concentration or vendor on the basal cytokine secretion profiles of differentiated macrophages (**Figure S1B-C**), albeit some donors showed increased secretion of CXCL10, IL-1RA and IL-23 at higher concentrations. Morphology and adherence were assessed using brightfield microscopy and little variation was observed across tested M-CSF conditions (**Figure S1D**). Based on this data, we used 25ng/mL M-CSF to generate M0 macrophages in subsequent experiments as it generated the most consistent responses across our tested donors.

To determine how activation conditions impact macrophage phenotype, human macrophages were exposed to several cytokine combinations to elicit M1-like and M2-like activation profiles. The range of concentrations used in this experiment was consistent with those reported previously in the literature (**Figure 1A and Table S1**). Overall, we tested eight total conditions for 3 male and 3 female donors. We first determined how these conditions impacted macrophage cytokine secretion profiles and found that variability in the data was mostly driven by polarization and not by concentration of stimulus (**Figure 1B and S2A**). Interestingly, one donor was characterized by elevated IL-1β and IL-6 in every condition compared to the other donors (**Figure S2B-C**).

**Figure 1.**
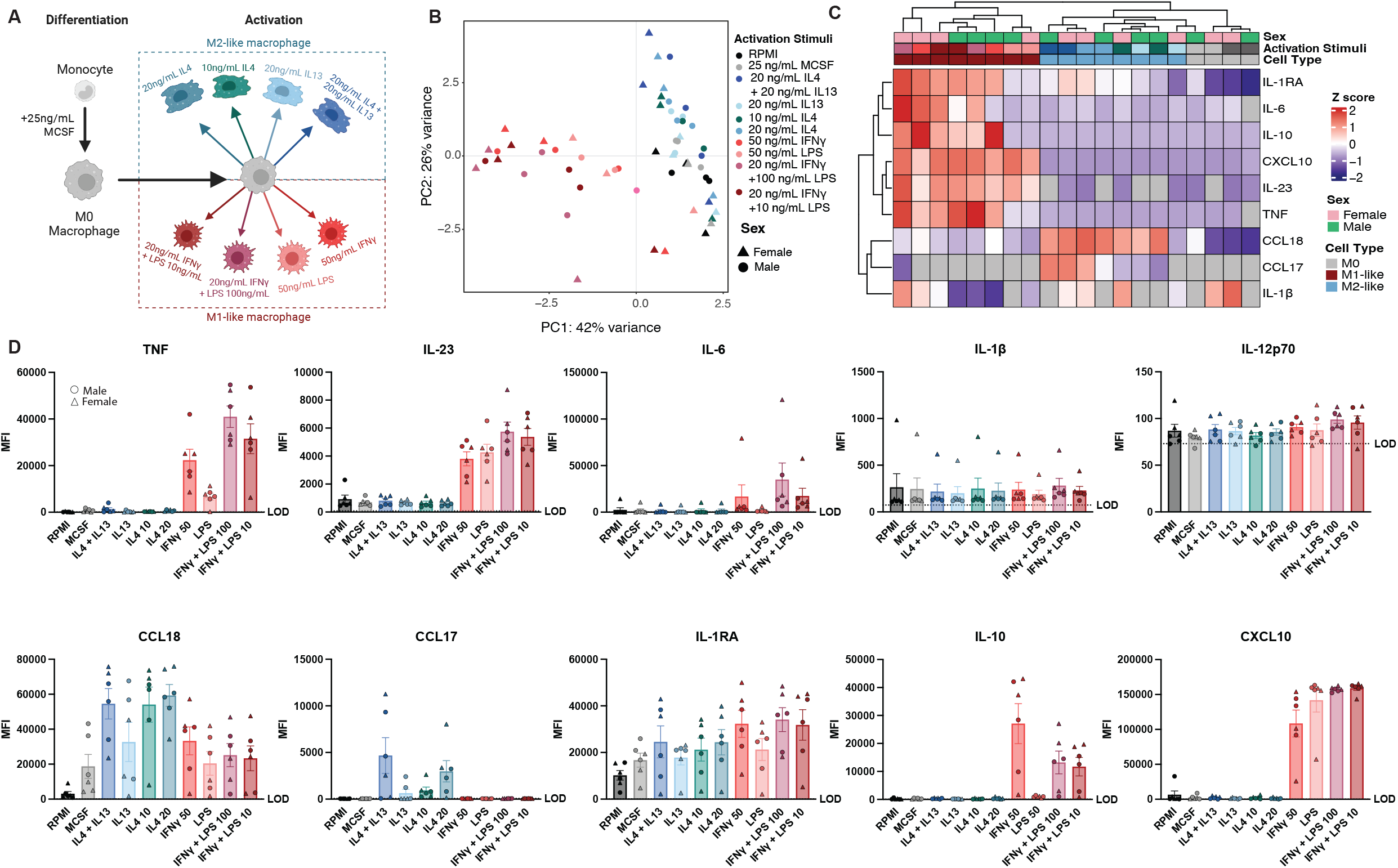
Macrophage activation stimuli generate cells with distinct cytokine secretion profiles. (A) Schematic of stimuli combinations and concentrations used for activating M0 macrophages into M1-like and M2-like macrophages. (B) Principal component analysis of macrophages treated with the respective activating stimuli (n=3) donors per sex. (C and D) Cytokines quantified in supernatants of differentiated M1-like and M2-like macrophages. Limit of (LOD) is 2 standard deviations above the average of the blank reading per cytokine. (C) Heatmap shows averaged mean fluorescence intensity (MFI) of 3 donors per condition. Data beneath the LOD is represented as a grey tile. Data is row normalized, and Z scored. (D) Bar plots of measured cytokines are shown. Male and female donors are indicated by circles or triangles, respectively. Bar is at the mean with the standard error of the mean (SEM) plotted.

Notably, there was variability in the magnitude of cytokine secretion within each polarization condition tested (**Figure 1C)**. For example, we found the highest level of IL-23 and IL-6 production when 20ng/mL of interferon gamma (IFNγ) was used with 100ng/mL of lipopolysaccharide (LPS), and the lowest overall when LPS alone was used for generating M1-like macrophages (**Figure 1D**). Similarly, we also observed variability across conditions for CCL18 and CCL17 for generating M2-like macrophages (**Figure 1E**). Finally, some cytokines, like IL-10 and CXCL10, showed higher production in M1-like macrophages across all conditions, while IL-1β and IL-12p70 levels were similar across conditions. Taken together, these results are consistent with prior reports demonstrating polarizing stimuli modulate macrophage immunophenotypes. Polarization with IFNγ and LPS at 10ng/mL leads to strong M1-like activation defined by production of TNF, IL-23, and IL-6, while polarization with IL-4 at 20ng/mL leads to robust M2-like activation characterized by CCL18 and IL-1RA production. Finally, we did not observe differences as a function of biological sex within each condition.

### Characterization of the transcriptome of un-stimulated M0 and activated M1-like and M2-like macrophages

Given these phenotypic differences, we next profiled the transcriptional states of M0, M1-like and M2-like macrophages to confirm their identity ^12,16,19^. Bulk RNA-sequencing was performed on a subset of the samples in Figure 1 to evaluate transcriptional differences between varying concentrations of LPS and M-CSF for M1-like and M0 macrophages, respectively, for two male and two female samples per group. These profiles were then compared to M2-like macrophages treated with 20ng/mL of IL-4 to identify the gene set profiles associated with each macrophage activation state. As observed in Figure 2A, each activation state had a distinct transcriptional profile, separating clearly along principal components 1 and 2. Interestingly, we did not see clear separation by biological sex. Using the DESeq2 package ^29^, we applied rlog transformation to the count data and identified the top 35 most variable genes by calculating gene-wise variance across all samples. Most of these genes were uniquely upregulated in M1-like macrophages, with relatively few genes shared between M0 and M2-like macrophages, highlighting distinct transcriptional profiles associated with macrophage polarization states. In M1-like macrophages, key inflammatory genes were identified, including: *CXCL9, CXCL10, CXCL11*, which are IFNγ-inducible and recruit Th1 cells; *GBP1* and *GBP5*, which have inflammasome activating roles; and inflammation-associated genes like *IDO1, IFIT3, IFITM1*, and *SERPING1* (**Figure 2B**). In M2-like macrophages, *F13A1, CCL18, CCL13* and *DNASE1L3* were upregulated; each of these is associated with cellular repair mechanisms, immunoregulation and resolution of inflammation. Expression of *ADAM19, CCR7, CCL5, C1S, CCL8, MT1G, MT2A*, and *MT1H* was common to both M1-like and M2-like macrophages, albeit at different expression levels. These genes show a gradient of expression with high levels in M1-like macrophages, moderate levels in M2-like macrophages, and low levels in M0 macrophages, reflecting their shared roles in macrophage activation. In M1-like cells, these genes drive pro-inflammatory responses, including immune cell recruitment (*CCL5, CCL8*), complement activation (*C1S*), and tissue remodeling (*ADAM19*). CCR7 supports migration to lymphoid tissues, while metallothionein genes (*MT1G, MT2A, MT1H*) mitigate oxidative stress during inflammation. In M2-like macrophages, gene expression aids in tissue repair and immune regulation, whereas M0 macrophages remain largely inactive with low gene expression.

**Figure 2.**
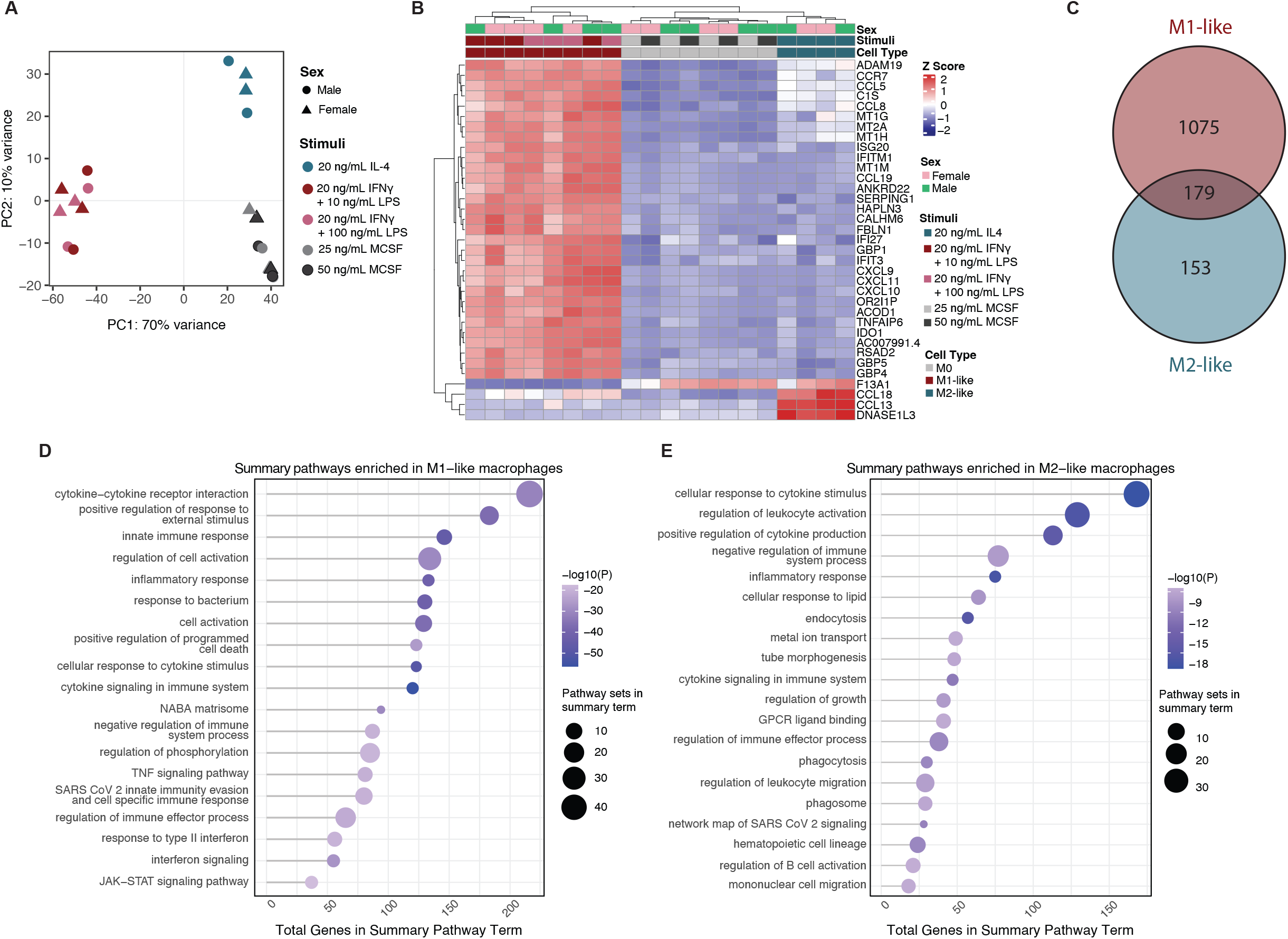
Activated macrophages reveal distinct transcriptional profiles. (A) Principal component analysis of macrophage transcriptomes. (B) Heatmap of 35 most variable genes within the dataset with biological sex, activation stimuli and cell type shown. (C) Venn diagram of differentially expressed genes in each activation state compared to M0 macrophages. (D and E) Pathway analysis results of M1-like macrophage DEG (D) or M2-like macrophages (E) compared to M0 in Metascape.

We next performed differential gene expression analysis to identify statistically significant genes that drive differences across each phenotype. When we compared the transcriptomes of M0 macrophages treated with 50ng/mL of M-CSF to those treated with 25ng/mL M-CSF, we saw no significant differentially expressed genes (DEGs) (**Figure S3A)**. We also saw no significant DEGs when we compared the transcriptomes of M1-like macrophages treated with 20ng/mL of IFNγ and 10ng/mL LPS vs 20ng/mL of IFNγ and 100ng/mL LPS (**Figure S3B)**. When we compared DEGs lists of M1-like and M2-like macrophages, we found that there were 1,075 unique to M1-like, 153 unique to M2-like and 179 DEGs that were shared between the two activation states (**Figure 2C and S3C-D**). To determine what biological functions were enriched in these gene lists that were unique to M1-like or M2-like macrophages, we performed pathway analysis. Pathways enriched in M1-like macrophages were generally related to their pro-inflammatory and immune-activating roles, and included cytokine-cytokine receptor interaction, innate immune response, inflammatory response, and cytokine signaling (**Figure 2D**). In contrast, the M2-like macrophages DEGs were enriched for pathways associated with immune regulation, tissue repair, and anti-inflammatory functions, and included cellular response to cytokine stimulus, regulation of leukocyte activation, and negative regulation of immune system process. Endocytosis, phagocytosis, and phagosome pathway enrichment suggests involvement in debris clearance and tissue homeostasis, while tube morphogenesis and regulation of growth emphasize roles in wound healing and angiogenesis. Enrichment in cytokine signaling and GPCR ligand binding further supports the role of M2-like macrophages in immune resolution and tissue remodeling characteristic of the M2-like phenotype (**Figure 2E)**. These transcriptional data, along with the cytokine secretion profiles, confirm that our optimized protocol reliably induces M1-like and M2-like macrophage activation states consistent with their well-characterized functional phenotypes.

### Rapid depolarization of cultured macrophages is observed once polarization media is removed

Given macrophage plasticity, we wanted to determine how quickly activated human macrophages lose their functional cytokine secretion profiles when activating stimuli within the media are removed^30–32^. Depolarization refers to macrophages transitioning from one state to another in response to environmental changes, like removal of activating stimuli. The timing of depolarization varies based on stimuli and conditions, but has been reported to range from several hours to several days in culture^6,33^. This has important implications for cell and tissue engineering, as well as for generating macrophages that can be used to study temporal processes *ex vivo*.

M1-like and M2-like macrophages were generated using the optimized protocol outlined in Figures 1 and 2 and activated for 24 hours. Activation stimuli were then either removed or maintained in culture for an additional 72 hours (**Figure 3A**). Strikingly, TNF, IL-6, and IL-1RA production decreased significantly after 24 hours of depolarization in both M1-like and M2-like macrophages. IL-1β secretion decreased in M1-like macrophages but remained constant for M2-like macrophages in the presence and absence of activating stimuli (**Figure 3B**). We found several cytokines, like IL-23 and CXCL10, however, remained largely unaffected (**Figure S4**). Taken together, this suggests that depolarization happens rapidly and dramatically impacts the cytokine profiles and likely the function of macrophages. This has implications for functional assays with human macrophages, as these data suggest that polarizing stimuli should be maintained in culture for the entire experimental duration.

**Figure 3.**
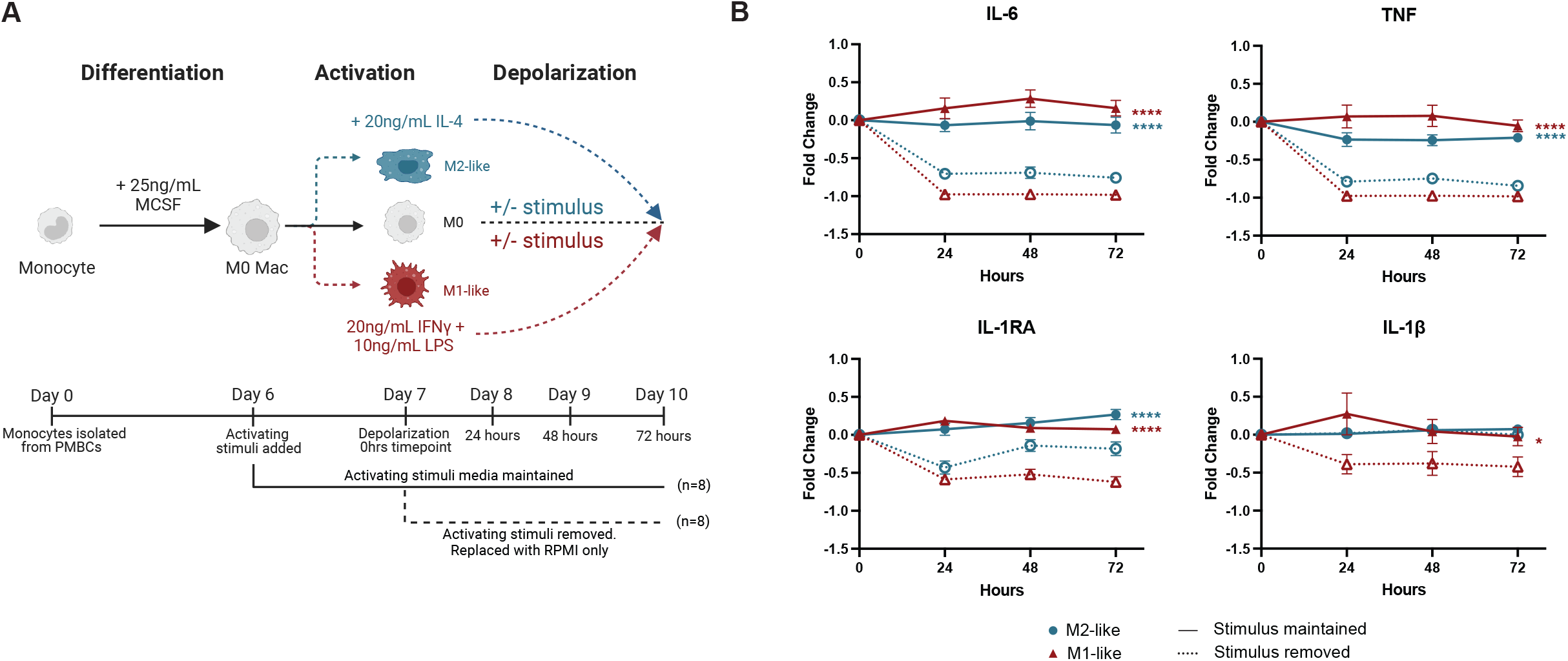
Removal of stimulus causes rapid depolarization of cultured macrophages. (A) Schematic of depolarization experiment. (B) Cytokine secretion of profiles of activated macrophages in the presence or absence of activating stimuli from 0 to 72 hours. Data is represented as fold change relative to the 0 hr timepoint. Significance is calculated using a two-way ANOVA with the interaction term of activating stimuli and time is plotted. *p < 0.05, ****p < 0.0001.

### Isolation of nuclei from cultured macrophages enables snRNA-seq workflows for streamlined profiling

Single-cell methods, like single-cell and single-nuclei RNA-sequencing (scRNA-seq and snRNA-seq), are essential for characterizing macrophage heterogeneity^34,35^. To mitigate concerns associated with mechanical and chemical dislodging of human macrophages from culture plates and subsequent effects on transcriptomics, we developed a workflow to isolate nuclei for snRNA-seq directly from culture plates to streamline profiling and minimally perturb the macrophage transcriptomen ^36,37^. We created an optimized isolation protocol that leverages 0.1% NP40 to isolate thousands of nuclei per culture well for snRNA-seq with 10X Genomics workflows (**Figure 4A**). We combined this with oligo-hashing to pool samples across many wells for sequencing, which is important for batch effect and cost considerations. This approach generated high quality snRNA-seq data with a high number of detected genes and a low percent of reads mapping to mitochondrial genes (**Figure S5A**). We also recovered all input pools using oligo tags (**Figure 4B** and **Figure S5B**), and filtered out all cells that didn’t have a single hashtag oligo and a unique molecular identifier (UMI). Cells with more than 5% mitochondrial gene expression were removed from analysis and we regressed out ribosomal genes, mitochondrial genes and donor (**Figure S5C-D**). As demonstrated in Figure 4C, macrophages show a remarkable degree of heterogeneity under 2D culture conditions. We identified four main clusters of cultured macrophages that have distinct transcriptional signatures and marker genes that clearly define their transcriptome but are not related to cell cycle (**Figure 4D**). To explore the functional roles of macrophage subpopulations identified by snRNA-seq, we performed pathway enrichment analysis using Metascape on cluster-specific marker genes identified through differential expression analysis. Markers were defined as genes with a log2 fold change greater than 1 and expressed in at least 60% of cells within a cluster. Cluster 0 is enriched for pathways related to receptor tyrosine kinase signaling, cytoskeletal remodeling, and cell differentiation, suggesting a role in signal transduction and cellular plasticity (**Figure S6A**). Cluster 1 shows strong enrichment for cytokine signaling, antigen presentation, and pathogen response, consistent with a pro-inflammatory, immune-activated macrophage state (**Figure S6B**). Cluster 2 is characterized by pathways involved in Fc receptor signaling, autophagy, and lysosomal function, indicative of a patrolling or homeostatic maintenance phenotype (**Figure S6C)**. In contrast, Cluster 3 is enriched for endocytosis, EGFR signaling, and suppression of immune activation, suggestive of a pro-reparative or immunomodulatory state (**Figure S6D)**. We also see that several canonical markers of macrophages either have very homogeneous or heterogeneous expression across all macrophages in our dataset (**Figure 4F**). Taken together, these data show that our approach can be used to profile macrophages with snRNA-seq in a cost effective and streamlined way.

**Figure 4.**
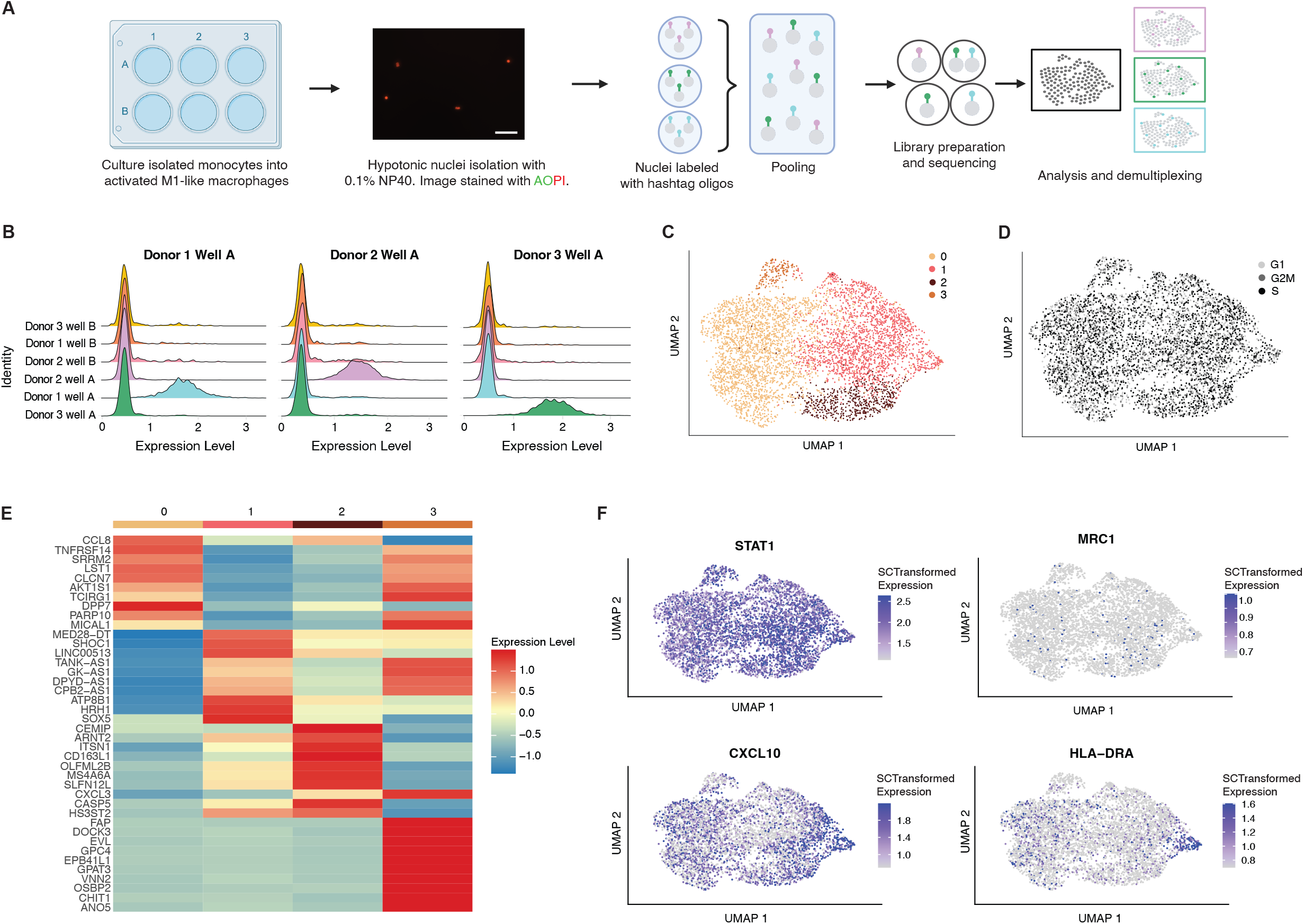
Characterization of macrophage heterogeneity with snRNA-seq. (A) Workflow schematic showing isolation of nuclei and library generation. (B) Representative plots of using hashing oligo demultiplexing for 3 donor pools. (C) UMAP of cells with 4 main clusters. (D) UMAP of cells colored by cell cycle score. (E) Heatmap of average gene expression of top 10 marker genes per cluster. (F) Feature plots of key marker genes.

## Discussion

We sought to systematically determine how variations in cytokine combinations, source vendor, and concentrations used impact the culture of human macrophages. Our study provides comprehensive and important insights into the factors influencing human macrophage activation and culture ex vivo and provides a robust culture system with validated reagents that are readily commercially available. We also used several methods to characterize macrophages after activation, including assaying cytokine secretion profiles and their transcriptional landscapes. Finally, we show that nuclei can be isolated readily from cultured macrophages for snRNA-seq, which is important for profiling these heterogeneous cells.

Protocols for culturing macrophages vary widely, including differences in CD14^+^monocyte isolation methods and the concentration or choice of M-CSF or GM-CSF used for differentiation. Prior studies have shown that GM-CSF induces inflammatory profiles in differentiated macrophages prior to activation^13,38,39^. Additionally, M-CSF concentration has been reported to influence macrophage differentiation, with high doses leading to impaired myeloid differentiation and a shift toward a dendritic cell-like phenotype^40^ as well as inducing variable morphological changes^41^. Here, we found that neither the concentration nor the source of M-CSF significantly affected the baseline phenotype of human monocyte-derived macrophages, suggesting that, within the tested range, M-CSF concentration does not substantially impact cytokine secretion or transcriptional activity.

We also determined how cocktails of immune stimuli impacted the activation of macrophages into M1-like or M2-like phenotypes. M1-like macrophages, typically activated by LPS and IFNγ, are well-documented to secrete pro-inflammatory cytokines such as TNF, IL-6, IL-1β, and CXCL10^11–14^. In contrast, M2-like macrophages, stimulated with IL-4 and/or IL-13, are associated with the production of CCL17, CCL18, and IL-10, which support tissue remodeling and anti-inflammatory responses^11,42,43^. Interestingly, we observed that IL-10, a cytokine traditionally linked to M2-like macrophages, was also produced under IFNγ stimulation. This suggests that anti-inflammatory feedback mechanisms may be engaged even in strongly pro-inflammatory environments, consistent with prior reports of context-dependent IL-10 induction^44,45^. Our data also supports the prevailing view that macrophage activation exists along a spectrum, rather than fitting into a strict M1/M2 dichotomy, as shown by PCA and clustering analyses revealing gradual transitions and overlapping cytokine profiles. The simultaneous secretion of pro- and anti-inflammatory cytokines—such as IL-23 and IL-10 under IFNγ + LPS stimulation—further highlights this plasticity and suggests that macrophages integrate multiple environmental signals to maintain adaptability. This variability even in monoculture presents challenges for generating cell-based macrophage therapies, and further investigation into factors that can better standardize cultures is warranted.

Our transcriptomic analysis revealed distinct gene expression signatures associated with M1-like and M2-like macrophages, which is consistent with prior reports. The enrichment of inflammatory pathways in M1-like macrophages and tissue repair pathways in M2-like macrophages aligns with their respective roles in immune responses. The gene sets that we identified for macrophages stimulated with LPS and IFNγ (for M1-like) and IL-4 (for M2-like) respectively align well with what has been previously reported for human cultured macrophages^11,12,16,33^. Additionally, we observed that a subset of genes was expressed across both M1-like and M2-like macrophage populations, suggesting transcriptional overlap for possible core functions despite polarized stimulation. Prior studies have similarly reported co-expression of genes traditionally associated with opposing activation programs, particularly under transitional or tissue-specific conditions^23,46–48^.These shared genes may reflect a core macrophage identity program or represent intermediate activation states shaped by the extracellular environment. Our data contribute to this evolving framework by highlighting that even under defined polarizing conditions, macrophages retain transcriptional flexibility.

One of the key findings of our study is the rapid depolarization of macrophages upon removal of activation stimuli, indicating that *in vitro* macrophages are highly plastic and can revert to a less activated or intermediate state within a short time frame despite strong initial cell programming. This observation is consistent with prior reports demonstrating that macrophage activation states are not fixed, but instead dynamic and reversible, particularly in response to changes in the extracellular environment^30–32,49^. Previous studies have reported varying timelines for macrophage depolarization, ranging from as long as 72 hours^31^ to as short as 6–24 hours^49,50^ and investigated how this impacts cellular repolarization to newly-introduced stimuli. We observed rapid functional depolarization, marked by a reduction secretion of TNF, IL-6, IL-1RA, and IL-1β, following the removal of activating stimuli. However, several cytokines continued to be produced— CXCL10, IL-23, and CCL18 in M1-like macrophages, and CCL17 and CCL18 in M2-like macrophages—suggesting residual functional memory. This supports the concept of macrophage memory, which may contribute to the incomplete repolarization of M1-like macrophages into M2-like states, as reported in previous studies that quantified cytokine secretion^49,51,52^ that was not observed in a transcriptional only investigation^31^. The implications of this finding have both experimental and translational relevance, as they highlight the importance of maintaining polarizing stimuli to preserve macrophage phenotype *in vitro*. Further, our results provide evidence for the inherent plasticity of macrophages and the potential for incomplete repolarization and raise important considerations for therapeutic strategies aiming to modulate macrophage function in disease.

Our adaptation of snRNA-seq for macrophages also provides a valuable tool for studying macrophage heterogeneity without the challenges associated with cell detachment. Prior work has shown that dissociation methods can significantly impact the resulting transcriptome^36,37^, and that isolation of nuclei or the use of cold enzymes can help to mitigate signatures of dead and dying cells. Given that macrophages are incredibly responsive, there is a clear need for methods that allow for profiling these cells from culture without invasive detachment methods. We show here that we can easily obtain high-quality snRNA-seq data, which coupled with hashing oligos for pooling, makes these experiments very cost effective. snRNA-seq data also demonstrate that there is heterogeneity across the culture, as we identified four clusters defined by unique gene expression programs. These include macrophage populations characterized by programs involved in cellular signal transduction and plasticity, inflammatory activation, patrolling or homeostatic maintenance, and tissue repair and immunomodulation. By preserving the native cellular state, snRNA-seq can minimize artifacts of dissociation and provide deeper insights into macrophage biology.

While we did not observe major sex-specific differences in macrophage activation or cytokine profiles among the donors used in this study, the influence of sex on macrophage biology remains an important consideration. Previous studies have demonstrated that sex hormones, particularly estrogen and testosterone, can modulate macrophage function, including polarization, cytokine production, and responsiveness to stimuli^28,53–56^. Although our experimental conditions did not reveal statistically significant sex-driven differences, this may be due to limited donor numbers. Future work should investigate these differences *in vitro* with higher numbers of donors. As sex-based immune variation becomes increasingly recognized as a critical variable in immunological research, future studies should be designed to more directly investigate how sex and hormonal milieu influence macrophage phenotype and function in both physiological and pathological contexts.

Overall, our study underscores the importance of standardizing macrophage culture conditions, considering donor variability, and leveraging new technologies to better characterize macrophage function. We present a streamlined protocol for generation and characterization of human macrophages from cryopreserved PBMCs, including the use of snRNA-seq. Importantly, our work suggests that depolarization is rapid and can have functional consequences for macrophages in culture. By refining *in vitro* models and improving reproducibility, we hope to advance our understanding of macrophage biology and their roles in human health and disease, while also enabling the precise manipulation of these cells’ *ex vivo* for therapeutic applications.

## MATERIALS AND METHODS

### M-CSF optimization of monocyte differentiation

Monocytes were isolated from cryopreserved PBMCs using Stem Cell’s CD14+ negative isolation kit from 3 female and 3 male donors (Table S6). PBMCs were thawed in RPMI with 0.05% DNase I. Monocytes were seeded at a density of 100,000 cells per well in a 96 well tissue culture treated plate with MCSF in RPMI with 10% FBS and 1% pen-strep for 6 days. MCSF concentration was optimized using concentrations 0, 10, 25, 50, and 100 ng/mL.

### Optimization of activating stimuli

Following optimization of 25ng/mL of M-CSF during the 6-day differentiation, 4 different inflammatory stimuli cocktails and 4 different anti-inflammatory cocktails were used to produce varied cell phenotypes for the same donors. For M1-like stimuli, macrophages were treated with 20ng/mL IFNγ and 10ng/mL LPS, 20ng/mL IFNγ and 100ng/mL LPS, 50ng/mL IFNγ, and 50ng/mL of LPS. For M2-like stimuli, cells were cultured in 20ng/mL IL-4, 10ng/mL IL-4, 20ng/mL IL-13 and 20ng/mL IL-4 + 20ng/mL IL-13. Complete RPMI containing 25ng/mL M-CSF was removed at day 6 and replaced with RPMI containing the respective activating cocktail and cultured for 24 hours.

### Cytokine analysis via Luminex

Supernatants (200 µl) were collected from 96 well plates containing 100,000 cells per well. Supernatants were analyzed neat and at 1:10 dilution using the Human Luminex Discovery Assay (cat. # LXSAHM-10) measuring CCL17, CCL18, CXCL10, IL-1RA, IL-1β, IL-6, IL-10, IL-23 and TNF. Cytokines were reported in mean fluorescent intensity on a Luminex Flexmap 3D instrument. Each dilution was run in technical duplicates and the average per duplicate pair is represented.

### Bulk RNA-seq

Libraries for RNA-seq were generated from cultured macrophages lysed in RLT buffer with 1% β-mercaptoethanol. RNA was purified from lysates using an Rneasy mini kits (Qiagen). Smart-Seq2 cDNA synthesis was performed using Superscript III reverse transcriptase (Thermo Fisher). Whole transcriptome amplified (WTA) product was purified with AMPure XP beads (Beckman Coulter), quantified with Quanit-iT PicoGreen dsDNA Assay (Thermo Fisher), normalized, and used for preparing paired-end sequencing libraries with Nextera XT (Illumina, #FC-131) using the manufacturer’s instructions. Libraries were pooled at 2M before sequencing on a NextSeq500 (Illumina) using a 75 cycle v2 sequencing kit.

### Analysis of transcriptomic data

After sequencing, BCL files were converted to FASTQs that were merged and demultiplexed. FASTQs were assessed for size to ensure that sequencing depth was comparable across all samples. They were then mapped to the GRCh38.p14 genome using STAR with count matrixes generated with HT-seq. Differential expression analysis was completed using the DESeq2 package in R.

### Macrophage Functional Depolarization

Monocytes were isolated from four female and four male donors and seeded at a density of 100,000 cells per well in a 96-well plate. Cells were cultured in the presence of M-CSF at a concentration of 25 ng/mL for six days at 37°C with 5% CO_2_. On Day 6, media were replaced with RPMI containing 20 ng/mL IFNγ and 10 ng/mL LPS for M1-like macrophage polarization, 20 ng/mL IL-4 for M2-like macrophages, or RPMI only for M0 macrophages. After 24 hours of stimulation, cells were either maintained in their respective media or washed and media was replaced with fresh RPMI in the absence of stimulation to assess functional depolarization. Supernatants were collected at 0-, 24-, 48-, and 72-hours post-stimulation and analyzed for cytokine secretion.

### Nuclei preparation for single nuclei sequencing with hashtag oligos

All steps were done at 4C, unless noted otherwise. Cultured cells were washed with PBS, then incubated on ice for 5min with 100ul hypotonic buffer (10mM Tris-HCl pH7.4, 10mM NaCl, 3mM MgCl2, 0.1% NP40). After 5min incubation, wells were loosely scraped with a pipette tip and 200ul of wash buffer (PBS 1X, 1% BSA, 0.2U/ul Protector RNAse Inhibitor) was added to the lysis solution in the well, pipette-mixed 5 times and transferred to a 1.5ml tube. An additional 500ul was used to wash the well and pipette-mixed with the transferred lysate an additional 5 to 10 times. Nuclei were spun at 600xg for 5min, then washed a second time before resuspending in 60ul of resuspension buffer (PBS 1X, 0.04%BSA, 0.2U/ul Protector RNAse Inhibitor) for nuclei counting and viability check using a Cellometer K2 automated cell counter (Revvity) and AO/PI live/dead reagent (Revvity). Nuclei integrity was also checked on EVOS M7000 (Life Technologies).

Nuclei tagging was done using the MULTIseq methodology ^57^in tandem with the Curiox washing system (Curiox Biosystems). In brief, anchor lipid-oligo and Barcoding oligos specific to each sample were incubated together then added to the respective samples to get incorporated into the nuclei membrane. A Co-anchor lipid-oligo was added to the mix to stabilize the tagging before transferring the samples to individual wells of a Curiox washing strip. Nuclei were incubated 40min on ice to bind oligos and allow nuclei to settle in the wells. Nuclei were washed 25x at flow rate of 1ul/s with PBS + 1% BSA prior to recovery, counting and pooling in equal proportions.

snSEQ was performed on the barcoded nuclei pool using the Chromium NextGem system (10X Genomics) and loaded targeting 20,000 nuclei. Gene expression and MULTIseq libraries were prepared and sequenced to a depth of 30,000 reads/cell on a NextSeq2000 instrument (Illumina) following manufacturer’s instructions.

### Analysis of single nuclei data

Cell Ranger was used for demultiplexing raw sequencing data and alignment. Initial quality filtering in Seurat v5^58^ (R 4.4.2) removed cells with >5% mitochondrial gene expression, fewer than 500 total expression counts, and a multiple or no hashtag oligos (HTO). The dataset was then normalized and scaled using SCTransform, regressing out mitochondrial genes, ribosomal genes, and HTO classification, followed by dimensionality reduction and clustering at a resolution of 0.2.

### Quantification and Statistical Analysis

Data are presented as mean ± standard error of the mean (SEM) unless specified differently. Statistical significance was analyzed using an ordinary one-way ANOVA for the activation and M-CSF experiments followed by Tukey’s correction for multiple comparisons. Depolarization experiments were analyzed using a two-way ANOVA or random mixed effects model was used to calculate the significance of the interaction term of activating stimuli over time. All statistical tests were generated in GraphPad Prism software (10.4.1). p values <0.05 were considered statistically significant indicated by *p < 0.05, **p < 0.01, ***p < 0.001, ****p < 0.0001. The number of donors included in each experiment is indicated in the figure legend.

## Acknowledgments

We would like the thank Dr. Margie Ackerman and Ackerman Lab members for helpful discussions. We would also like to thank Dr. Jonathan DeLong and Dr. Emily Morris for helpful discussions about macrophage detachment. This work was supported by Dartmouth Cancer Center Support Grant (P30CA023108/CA/NCI/NIH/HHS) and the Genomics & Molecular Biology (SCR_021293) Shared Resources (SCR_019165). B.A.G and O.G.P. are supported in part through the Geisel School of Medicine at Dartmouth’s Center for Quantitative Biology through a grant from the National Institute of General Medical Sciences (NIGMS, P20GM130454) of the NIH.

## Author Contributions

B.A.G. and O.G.P conceptualized the work, designed experiments, generated figures, and wrote the manuscript. O.G.P, L.P., and F.W.K collected data. O.G.P analyzed data.

P.P. contributed to experimental design, data interpretation, and manuscript writing. All authors participated in discussions regarding this manuscript and approved its final submission.

## REFERENCES

1. Park, M.D., Silvin, A., Ginhoux, F., and Merad, M. (2022). Macrophages in health and disease. Cell 185, 4259–4279. 10.1016/j.cell.2022.10.007.

2. Wynn, T.A., Chawla, A., and Pollard, J.W. (2013). Macrophage biology in development, homeostasis and disease. Nature 496, 445–455. 10.1038/nature12034.1.

3. Mosser, D.M., Hamidzadeh, K., and Goncalves, R. (2021). Macrophages and the maintenance of homeostasis. Cell. Mol. Immunol. 18, 579–587. 10.1038/s41423-020-00541-3.

4. Chen, S., Saeed, A.F.U.H., Liu, Q., Jiang, Q., Xu, H., Xiao, G.G., Rao, L., and Duo, Y. (2023). Macrophages in immunoregulation and therapeutics. Signal Transduct. Target. Ther. 8, 207. 10.1038/s41392-023-01452-1.

5. Gurvich, O.L., Puttonen, K.A., Bailey, A., Kailaanmäki, A., Skirdenko, V., Sivonen, M., Pietikäinen, S., Parker, N.R., Ylä-Herttuala, S., and Kekarainen, T. (2020). Transcriptomics uncovers substantial variability associated with alterations in manufacturing processes of macrophage cell therapy products. Sci. Rep. 10, 14049. 10.1038/s41598-020-70967-2.

6. Zheng, Y., Wei, K., Jiang, P., Zhao, J., Shan, Y., Shi, Y., Zhao, F., Chang, C., Li, Y., Zhou, M., et al. (2024). Macrophage polarization in rheumatoid arthritis: signaling pathways, metabolic reprogramming, and crosstalk with synovial fibroblasts. Front. Immunol. 15, 1394108. 10.3389/fimmu.2024.1394108.

7. Dussold, C., Zilinger, K., Turunen, J., Heimberger, A.B., and Miska, J. (2024). Modulation of macrophage metabolism as an emerging immunotherapy strategy for cancer. J. Clin. Investig. 134, e175445. 10.1172/jci175445.

8. Mills, C.D., Kincaid, K., Alt, J.M., Heilman, M.J., and Hill, A.M. (2000). M-1/M-2 Macrophages and the Th1/Th2 Paradigm. J. Immunol. 164, 6166–6173. 10.4049/jimmunol.164.12.6166.

9. Liu, J., Geng, X., Hou, J., and Wu, G. (2021). New insights into M1/M2 macrophages: key modulators in cancer progression. Cancer Cell Int. 21, 389. 10.1186/s12935-021-02089-2.

10. Peng, Y., Zhou, M., Yang, H., Qu, R., Qiu, Y., Hao, J., Bi, H., and Guo, D. (2023). Regulatory Mechanism of M1/M2 Macrophage Polarization in the Development of Autoimmune Diseases. Mediat. Inflamm. 2023, 8821610. 10.1155/2023/8821610.

11. Edwards, J.P., Zhang, X., Frauwirth, K.A., and Mosser, D.M. (2006). Biochemical and functional characterization of three activated macrophage populations. J. Leukoc. Biol. 80, 1298–1307. 10.1189/jlb.0406249.

12. Xue, J., Schmidt, S.V., Sander, J., Draffehn, A., Krebs, W., Quester, I., De Nardo, D., Gohel, T.D., Emde, M., Schmidleithner, L., et al. (2014). Transcriptome-Based Network Analysis Reveals a Spectrum Model of Human Macrophage Activation. Immunity 40, 274–288. 10.1016/j.immuni.2014.01.006.1.

13. Fleetwood, A.J., Dinh, H., Cook, A.D., Hertzog, P.J., and Hamilton, J.A. (2009). GM-CSF- and M-CSF-dependent macrophage phenotypes display differential dependence on Type I interferon signaling. J. Leukoc. Biol. 86, 411–421. 10.1189/jlb.1108702.

14. Gundra, U.M., Girgis, N.M., Ruckerl, D., Jenkins, S., Ward, L.N., Kurtz, Z.D., Wiens, K.E., Tang, M.S., Basu-Roy, U., Mansukhani, A., et al. (2014). Alternatively activated macrophages derived from monocytes and tissue macrophages are phenotypically and functionally distinct. Blood 123, e110–e122. 10.1182/blood-2013-08-520619.

15. Rutschman, R., Lang, R., Hesse, M., Ihle, J.N., Wynn, T.A., and Murray, P.J. (2001). Cutting Edge: Stat6-Dependent Substrate Depletion Regulates Nitric Oxide Production. J. Immunol. 166, 2173–2177. 10.4049/jimmunol.166.4.2173.

16. Murray, P.J., Allen, J.E., Biswas, S.K., Fisher, E.A., Gilroy, D.W., Goerdt, S., Gordon, S., Hamilton, J.A., Ivashkiv, L.B., Lawrence, T., et al. (2014). Macrophage Activation and Polarization: Nomenclature and Experimental Guidelines. Immunity 41, 14–20. 10.1016/j.immuni.2014.06.008.

17. Shirey, K.A., Cole, L.E., Keegan, A.D., and Vogel, S.N. (2008). Francisella tularensis Live Vaccine Strain Induces Macrophage Alternative Activation as a Survival Mechanism. J. Immunol. 181, 4159–4167. 10.4049/jimmunol.181.6.4159.

18. Lang, R., Patel, D., Morris, J.J., Rutschman, R.L., and Murray, P.J. (2002). Shaping Gene Expression in Activated and Resting Primary Macrophages by IL-10. J. Immunol. 169, 2253–2263. 10.4049/jimmunol.169.5.2253.

19. Lawrence, T., and Natoli, G. (2011). Transcriptional regulation of macrophage polarization: enabling diversity with identity. Nat. Rev. Immunol. 11, 750–761. 10.1038/nri3088.

20. Brennan, P.N., MacMillan, M., Manship, T., Moroni, F., Glover, A., Troland, D., MacPherson, I., Graham, C., Aird, R., Semple, S.I.K., et al. (2025). Autologous macrophage therapy for liver cirrhosis: a phase 2 open-label randomized controlled trial. Nat. Med. 31, 979–987. 10.1038/s41591-024-03406-8.

21. Reiss, K.A., Angelos, M.G., Dees, E.C., Yuan, Y., Ueno, N.T., Pohlmann, P.R., Johnson, M.L., Chao, J., Shestova, O., Serody, J.S., et al. (2025). CAR-macrophage therapy for HER2-overexpressing advanced solid tumors: a phase 1 trial. Nat. Med., 1– 12. 10.1038/s41591-025-03495-z.

22. Cess, C.G., and Finley, S.D. (2020). Multi-scale modeling of macrophage—T cell interactions within the tumor microenvironment. PLoS Comput. Biol. 16, e1008519. 10.1371/journal.pcbi.1008519.1.

23. Orecchioni, M., Ghosheh, Y., Pramod, A.B., and Ley, K. (2019). Macrophage Polarization: Different Gene Signatures in M1(LPS+) vs. Classically and M2(LPS–) vs. Alternatively Activated Macrophages. Front. Immunol. 10, 1084. 10.3389/fimmu.2019.01084.

24. Sica, A., and Mantovani, A. (2012). Macrophage plasticity and polarization: in vivo veritas. J. Clin. Investig. 122, 787–795. 10.1172/jci59643.

25. Bencheikh, L., Diop, M.K., Rivière, J., Imanci, A., Pierron, G., Souquere, S., Naimo, A., Morabito, M., Dussiot, M., Leeuw, F.D., et al. (2019). Dynamic gene regulation by nuclear colony-stimulating factor 1 receptor in human monocytes and macrophages. Nat. Commun. 10, 1935. 10.1038/s41467-019-09970-9.

26. Hou, X.-X., Wang, X.-Q., Zhou, W.-J., and Li, D.-J. (2021). Regulatory T cells induce polarization of pro-repair macrophages by secreting sFGL2 into the endometriotic milieu. Commun. Biol. 4, 499. 10.1038/s42003-021-02018-z.

27. Domschke, G., Linden, F., Pawig, L., Hafner, A., Akhavanpoor, M., Reymann, J., Doesch, A.O., Erbel, C., Weber, C., Katus, H.A., et al. (2018). Systematic RNA-interference in primary human monocyte-derived macrophages: A high-throughput platform to study foam cell formation. Sci. Rep. 8, 10516. 10.1038/s41598-018-28790-3.

28. Enright, S., and Werstuck, G.H. (2024). Investigating the Effects of Sex Hormones on Macrophage Polarization. Int. J. Mol. Sci. 25, 951. 10.3390/ijms25020951.

29. Love, M.I., Huber, W., and Anders, S. (2014). Moderated estimation of fold change and dispersion for RNA-seq data with DESeq2. Genome Biol. 15, 550. 10.1186/s13059-014-0550-8.

30. Oyarce, C., Vizcaino-Castro, A., Chen, S., Boerma, A., and Daemen, T. (2021). Re-polarization of immunosuppressive macrophages to tumor-cytotoxic macrophages by repurposed metabolic drugs. OncoImmunology 10, 1898753. 10.1080/2162402x.2021.1898753.

31. Liu, S.X., Gustafson, H.H., Jackson, D.L., Pun, S.H., and Trapnell, C. (2020). Trajectory analysis quantifies transcriptional plasticity during macrophage polarization. Sci. Rep. 10, 12273. 10.1038/s41598-020-68766-w.

32. Gharib, S.A., McMahan, R.S., Eddy, W.E., Long, M.E., Parks, W.C., Aitken, M.L., and Manicone, A.M. (2019). Transcriptional and functional diversity of human macrophage repolarization. J. Allergy Clin. Immunol. 143, 1536–1548. 10.1016/j.jaci.2018.10.046.

33. Hu, G., Su, Y., Kang, B.H., Fan, Z., Dong, T., Brown, D.R., Cheah, J., Wittrup, K.D., and Chen, J. (2021). High-throughput phenotypic screen and transcriptional analysis 1. identify new compounds and targets for macrophage reprogramming. Nat. Commun. 12, 773. 10.1038/s41467-021-21066-x.

34. Song, Y., Zhang, Q., Ban, R., Zhao, X., Sun, H., Lin, J., Guo, T., Wang, T., Xia, K., Xin, Z., et al. (2024). Single-nucleus RNA sequencing reveals that macrophages and smooth muscle cells promote carotid atherosclerosis progression through mitochondrial autophagy. Medicine 103, e37171. 10.1097/md.0000000000037171.

35. Lantz, C., Radmanesh, B., Liu, E., Thorp, E.B., and Lin, J. (2020). Single-cell RNA sequencing uncovers heterogenous transcriptional signatures in macrophages during efferocytosis. Sci. Rep. 10, 14333. 10.1038/s41598-020-70353-y.

36. Chen, S., So, E.C., Strome, S.E., and Zhang, X. (2015). Impact of Detachment Methods on M2 Macrophage Phenotype and Function. J. Immunol. Methods 426, 56–61. 10.1016/j.jim.2015.08.001.

37. Song, Q., Zhang, Y., Zhou, M., Xu, Y., Zhang, Q., Wu, L., Liu, S., Zhang, M., Zhang, L., Wu, Z., et al. (2022). The Culture Dish Surface Influences the Phenotype and Dissociation Strategy in Distinct Mouse Macrophage Populations. Front. Immunol. 13, 920232. 10.3389/fimmu.2022.920232.

38. Lotfi, N., Zhang, G.-X., Esmaeil, N., and Rostami, A. (2020). Evaluation of the effect of GM-CSF blocking on the phenotype and function of human monocytes. Sci. Rep. 10, 1567. 10.1038/s41598-020-58131-2.

39. Jiemy, W.F., Sleen, Y.van, Geest, K.S. van der, Berge, H.A.ten, Abdulahad, W.H., Sandovici, M., Boots, A.M., Heeringa, P., and Brouwer, E. (2020). Distinct macrophage phenotypes skewed by local granulocyte macrophage colony-stimulating factor (GM-CSF) and macrophage colony-stimulating factor (M-CSF) are associated with tissue destruction and intimal hyperplasia in giant cell arteritis. Clin. Transl. Immunol. 9, e1164. 10.1002/cti2.1164.

40. Menetrier-Caux, C., Montmain, G., Dieu, M.C., Bain, C., Favrot, M.C., Caux, C., and Blay, J.Y. (1998). Inhibition of the differentiation of dendritic cells from CD34(+) progenitors by tumor cells: role of interleukin-6 and macrophage colony-stimulating factor. Blood 92, 4778–4791.

41. Asakura, E., Hanamura, T., Umemura, A., Yada, K., Yamauchi, T., and Tanabe, T. (1996). Effects of Macrophage Colony-Stimulating Factor (M-CSF) on Lipopolysaccharide (LPS)-induced Mediator Production from Monocytes in vitro. Immunobiology 195, 300–313. 10.1016/s0171-2985(96)80047-7.

42. Gordon, S., and Martinez, F.O. (2010). Alternative Activation of Macrophages: Mechanism and Functions. Immunity 32, 593–604. 10.1016/j.immuni.2010.05.007.1.

43. Mantovani, A., Sica, A., Sozzani, S., Allavena, P., Vecchi, A., and Locati, M. (2004). The chemokine system in diverse forms of macrophage activation and polarization. Trends Immunol. 25, 677–686. 10.1016/j.it.2004.09.015.

44. Chang, E.Y., Guo, B., Doyle, S.E., and Cheng, G. (2007). Cutting Edge: Involvement of the Type I IFN Production and Signaling Pathway in Lipopolysaccharide-Induced IL-10 Production. J. Immunol. 178, 6705–6709. 10.4049/jimmunol.178.11.6705.

45. Su, X., Yu, Y., Zhong, Y., Giannopoulou, E.G., Hu, X., Liu, H., Cross, J.R., Rätsch, G., Rice, C.M., and Ivashkiv, L.B. (2015). Interferon-γ regulates cellular metabolism and mRNA translation to potentiate macrophage activation. Nat. Immunol. 16, 838–849. 10.1038/ni.3205.

46. Buscher, K., Ehinger, E., Gupta, P., Pramod, A.B., Wolf, D., Tweet, G., Pan, C., Mills, C.D., Lusis, A.J., and Ley, K. (2017). Natural variation of macrophage activation as disease-relevant phenotype predictive of inflammation and cancer survival. Nat. Commun. 8, 16041. 10.1038/ncomms16041.

47. Sanin, D.E., Ge, Y., Marinkovic, E., Kabat, A.M., Castoldi, A., Caputa, G., Grzes, K.M., Curtis, J.D., Thompson, E.A., Willenborg, S., et al. (2022). A common framework of monocyte-derived macrophage activation. Sci. Immunol. 7, eabl7482. 10.1126/sciimmunol.abl7482.

48. Jablonski, K.A., Amici, S.A., Webb, L.M., Ruiz-Rosado, J. de D., Popovich, P.G., Partida-Sanchez, S., and Guerau-de-Arellano, M. (2015). Novel Markers to Delineate Murine M1 and M2 Macrophages. PLoS ONE 10, e0145342. 10.1371/journal.pone.0145342.49.

49. Van den Bossche, J., Baardman, J., Otto, N.A., van der Velden, S., Neele, A.E., van den Berg, S.M., Luque-Martin, R., Chen, H.-J., Boshuizen, M.C.S., Ahmed, M., et al. (2016). Mitochondrial Dysfunction Prevents Repolarization of Inflammatory Macrophages. Cell Rep. 17, 684–696. 10.1016/j.celrep.2016.09.008.

50. Rückerl, D., Jenkins, S.J., Laqtom, N.N., Gallagher, I.J., Sutherland, T.E., Duncan, S., Buck, A.H., and Allen, J.E. (2012). Induction of IL-4Rα–dependent microRNAs identifies PI3K/Akt signaling as essential for IL-4–driven murine macrophage proliferation in vivo. Blood 120, 2307–2316. 10.1182/blood-2012-02-408252.

51. Davis, M.J., Tsang, T.M., Qiu, Y., Dayrit, J.K., Freij, J.B., Huffnagle, G.B., and Olszewski, M.A. (2013). Macrophage M1/M2 Polarization Dynamically Adapts to Changes in Cytokine Microenvironments in Cryptococcus neoformans Infection. mBio 4, 10.1128/mbio.00264-13. 10.1128/mbio.00264-13.

52. Khallou-Laschet, J., Varthaman, A., Fornasa, G., Compain, C., Gaston, A.-T., Clement, M., Dussiot, M., Levillain, O., Graff-Dubois, S., Nicoletti, A., et al. (2010). Macrophage Plasticity in Experimental Atherosclerosis. PLoS ONE 5, e8852. 10.1371/journal.pone.0008852.50.

53. Villa, A., Vegeto, E., Poletti, A., and Maggi, A. (2016). Estrogens, Neuroinflammation, and Neurodegeneration. Endocr. Rev. 37, 372–402. 10.1210/er.2016-1007.

54. Chen, K.-H.E., Lainez, N.M., and Coss, D. (2021). Sex Differences in Macrophage Responses to Obesity-Mediated Changes Determine Migratory and Inflammatory Traits. J. Immunol. 206, 141–153. 10.4049/jimmunol.2000490.

55. Keselman, A., Fang, X., White, P.B., and Heller, N.M. (2017). Estrogen Signaling Contributes to Sex Differences in Macrophage Polarization during Asthma. J. Immunol. 199, 1573–1583. 10.4049/jimmunol.1601975.

56. Becerra-Díaz, M., Lerner, A.D., Yu, D.H., Thiboutot, J.P., Liu, M.C., Yarmus, L.B., Bose, S., and Heller, N.M. (2021). Sex differences in M2 polarization, chemokine and IL-4 receptors in monocytes and macrophages from asthmatics. Cell. Immunol. 360, 104252. 10.1016/j.cellimm.2020.104252.

57. McGinnis, C.S., Patterson, D.M., Winkler, J., Conrad, D.N., Hein, M.Y., Srivastava, V., Hu, J.L., Murrow, L.M., Weissman, J.S., Werb, Z., et al. (2019). MULTI-seq: sample multiplexing for single-cell RNA sequencing using lipid-tagged indices. Nat. Methods 16, 619–626. 10.1038/s41592-019-0433-8.

58. Butler, A., Hoffman, P., Smibert, P., Papalexi, E., and Satija, R. (2018). Integrating single-cell transcriptomic data across different conditions, technologies, and species. Nat. Biotechnol. 36, 411–420. 10.1038/nbt.4096.

