## Supplemental Figures for "A streamlined and comprehensive protocol for the generation and multi-omic analysis of human monocyte-derived macrophages"

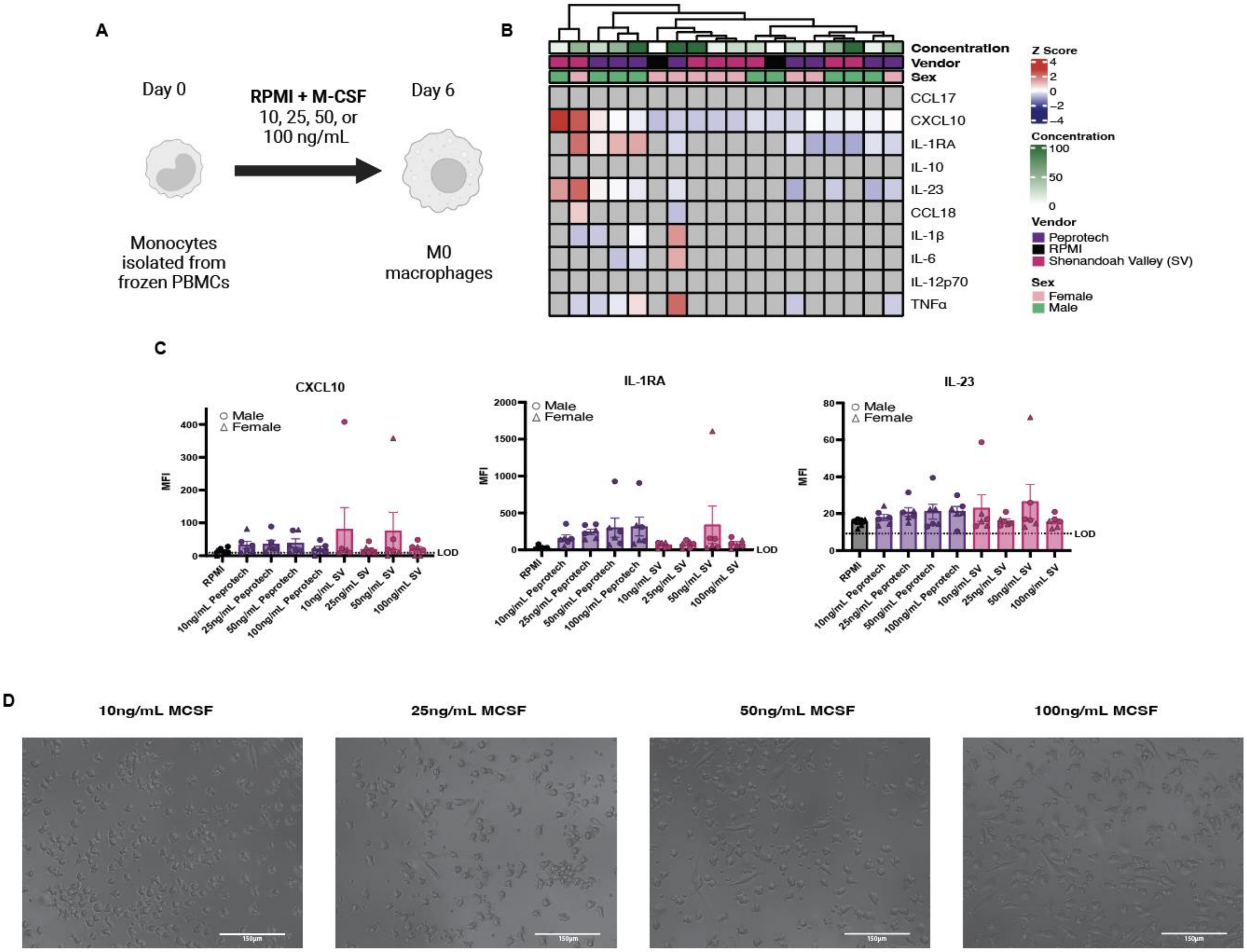


**Supplemental Figure 1: M-CSF Differentiation yields functionally active macrophages**
(A) Schematic of monocyte to macrophage differentiation. (B and C) Cytokines quantified in supernatants of differentiated M0 macrophages. Limit of (LOD) is 2 standard deviations above the average of the blank readings per cytokine. (B) Heatmap shows averaged mean fluorescence intensity (MFI) of 3 donors per condition that is above the limit of detection (LOD). Data beneath the LOD is represented as a grey tile. Data is row normalized, and Z scored. (C) Bar plots of measured cytokines are shown. Male and female donors are indicated by circles or triangles, respectively. Bar is at mean with SEM plotted. (D) Representative brightfield microscopy images of cultured macrophages at 120 hours.

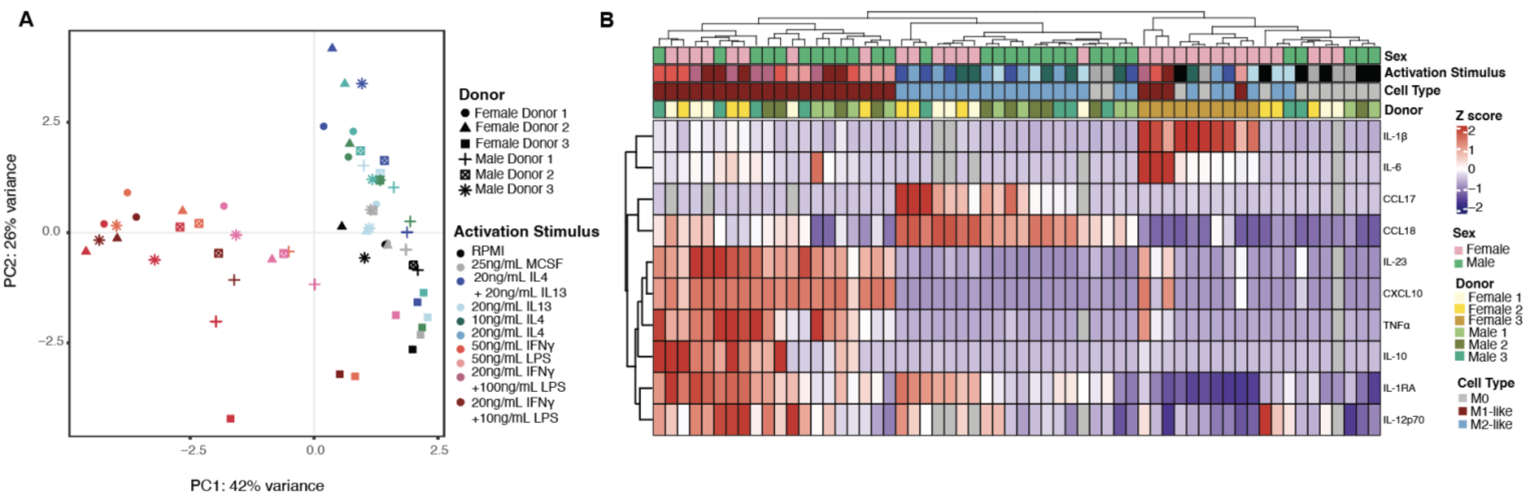


**Supplemental Figure 2: Donor variability within macrophage activation** (A) Principal component analysis of macrophages treated with the respective activating stimuli. (B) Cytokines quantified in supernatants of differentiated M1-like and M2-like macrophages. Limit of (LOD) is 2 standard deviations above the average of the blank reading per cytokine. Heatmap shows averaged mean fluorescence intensity (MFI) of 2 technical replicates. Data beneath the LOD is represented as a grey tile. Data is row normalized, and Z scored.

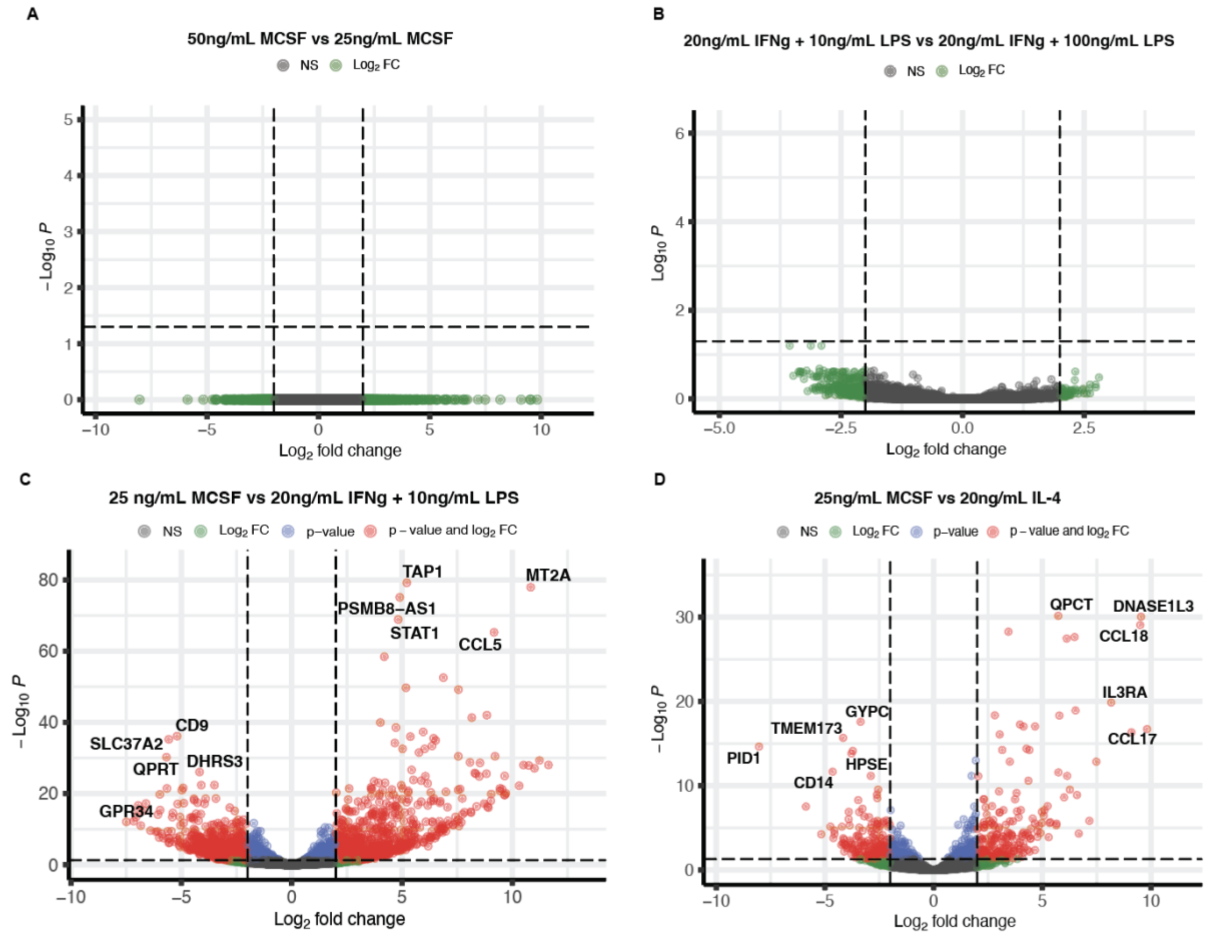


**Supplemental Figure 3: Transcriptional variation of activating stimuli with varied concentrations** (A-D) Volcano plots of differentially expressed genes of macrophages treated with variant concentrations of (n=4). (A) M0 macrophages treated with 25ng/mL or 50ng/mL of M-CSF 2 female and 2 male donors. (B) M1-like macrophages treated with 20ng/mL IFNγ and 10ng/mL LPS or 20ng/mL IFNγ and 100ng/mL LPS. (C) M1-like macrophages treated with 20ng/mL IFNγ and 10ng/mL LPS vs 25ng/mL M-CSF. (D) M2-like macrophages treated with 20ng/mL IL-4 vs 25ng/mL M-CSF.

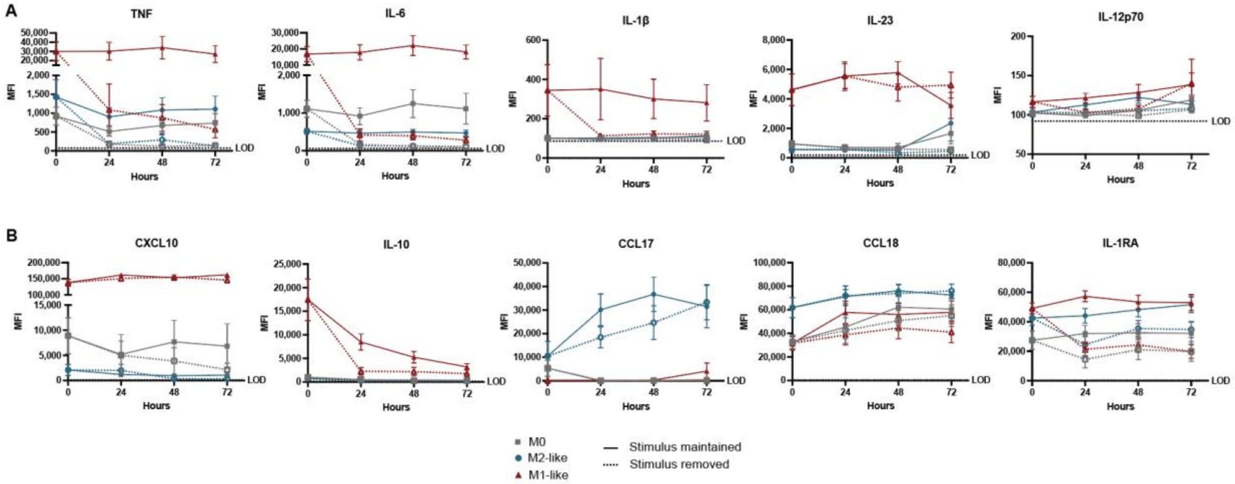


**Supplemental Figure 4: Depolarization of cultured macrophages.** (A-B) Cytokine secretion of profiles of activated macrophages in the presence or absence of activating stimuli from 0 to 72 hours (n=8).

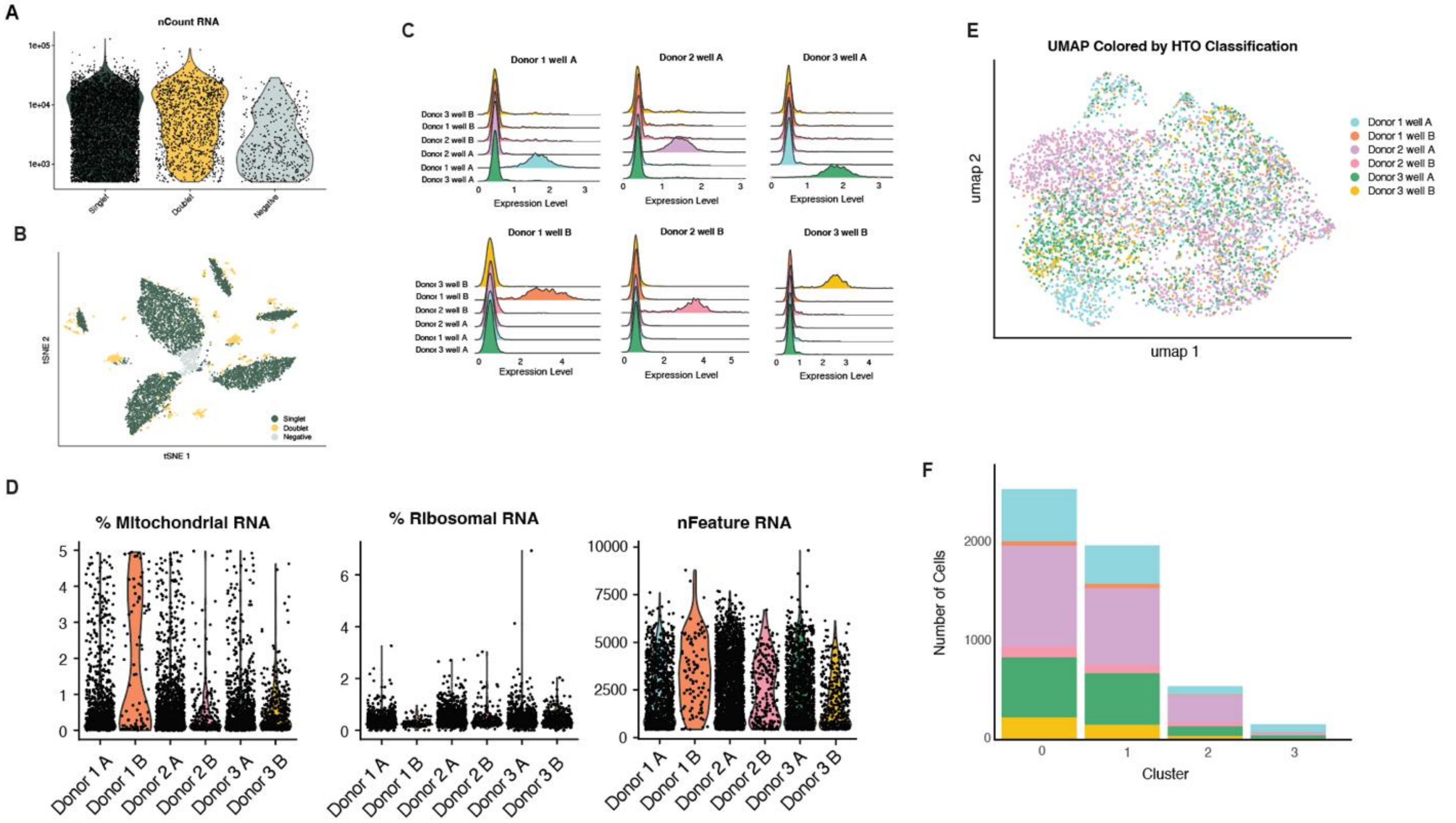


**Supplemental Figure 5: Quality statistics of snRNA-seq. (**A) Violin plots of hashtag oligo detection per cell. (B) tSNE of HTOs. (C) Ridge plots of hashtag oligo demultiplexing for all 6 wells. (D) Violin plots of RNA quality statistics split by HTO classification. (E) RNA umap colored by HTO donor. (F) Stacked bar plot of cell count per donor in each cluster.

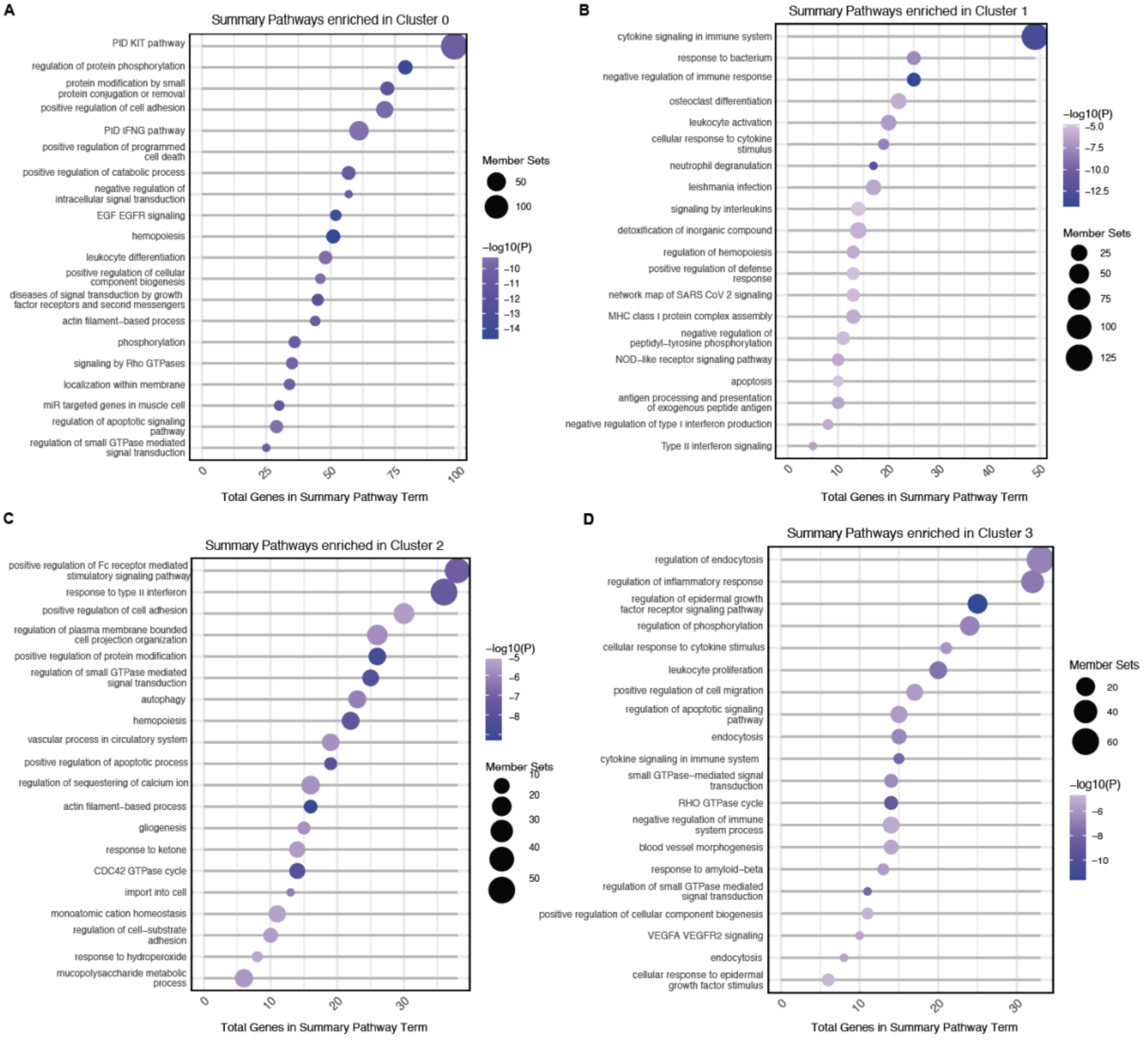


**Supplemental Figure 6: Pathway results of snRNA-seq clusters.** (A-D) Pathway analysis of clusters within M1-like macrophages in Metascape. (A) Cluster 0, (B) Cluster 2, (C) Cluster 3, (D) Cluster 4.
